# Ribosomal stress-induced senescence as a novel pro-senescence strategy for p16 positive basal-like breast cancer

**DOI:** 10.1101/469445

**Authors:** Madeleine Moore, Luke Gammon, Sally Dreger, James Koh, James C Garbe, Martha R Stampfer, Michael Philpott, Louise Jones, Cleo L Bishop

**Affiliations:** CBCR, The Blizard Institute, Barts and The London School of Medicine and Dentistry, Queen Mary University of London, 4 Newark Street, London E1 2AT, UK; Barts Cancer Institute, Barts and The London School of Medicine and Dentistry, Queen Mary University of London, 4 Newark Street, London E1 2AT, UK; Division of Surgical Sciences, Department of Surgery, Duke University Medical School, Durham, NC, 27710, USA.; Biological Systems & Engineering Division, Lawrence Berkeley National Laboratory, Berkeley, CA, 94720, USA.

## Abstract

Re-engaging the senescent programme represents an attractive yet underexplored strategy for cancer therapy, particularly for those tumour subtypes where targeted agents are limited or unavailable. Here, we identify a specific subset of ribosomal proteins as novel pro-senescence therapeutic targets for a highly aggressive subtype of breast cancer, p16 positive basal-like breast cancer. Mechanistically, ribosomal stress-induced senescence generates a stable cell cycle arrest, is dependent on endogenous p16 triggering a resensitisation to the p16/RB tumour suppressor axis, followed by establishment of a senescence-associated secretory phenotype, and is independent of DNA damage. Conversely, ribosomal protein knockdown in a p16 negative breast cancer model results in caspase-mediated apoptosis. Importantly, individual ribosomal protein loss is well tolerated by a panel of normal human cells. We demonstrate a reciprocal feedback loop between loss of RPS3A and RPS7 at both the transcriptional and post-transcriptional level during ribosomal stress-induced senescence. Further, our ribosomal hits are co-ordinately dysregulated in breast cancer, with elevated expression associated with a poor prognosis. Clinical relevance is demonstrated in tissue microarrays, and a RPS3A^HIGH^RPS7^HIGH^ signature is associated with an earlier disease onset and synergises with p16 to further worsen patient outcome. We conclude that dysregulation of ribosomal proteins constitutes a cancer cell-specific mechanism of senescence evasion and that engaging ribosomal stress-induced senescence may be relevant for future pro-senescence therapies.

## INTRODUCTION

Basal-like breast cancers (BLBCs) are an aggressive, early onset tumour type, commonly found to be triple negative (HER2, ER and PR negative) ^1^, and intriguingly are frequently positive for the tumour suppressor and senescence biomarker, p16^INK4a^ (p16) ^2^ whilst lacking retinoblastoma protein (RB). p16 positive BLBCs are often unresponsive to endocrine therapies or targeted agents, such as trastuzumab, frequently leaving radiation/chemotherapy as the only treatment option. p16 expression is associated with a particularly aggressive phenotype together with a poorer prognosis, and as such there is a pressing unmet clinical need for novel therapeutic targets for p16 positive BLBC.

The application of specific chemotherapeutics frequently activates a cancer-cell senescence programme (termed therapy-induced senescence, TIS) in a wide array of tumours, including breast cancer ^3^. Several studies show that cancer cell senescence can be activated via p53 restoration, telomerase inhibition or CDK modulation ^3^, supporting targeted pro-senescence anti-cancer strategies ^4^. The concept of activating this cellular programme for therapeutic gain in the most clinically challenging cancer subtypes is gaining momentum for two main reasons. Firstly, senescent cells secrete a panel of pro-inflammatory cytokines, termed the senescence-associated secretory phenotype (SASP), such as interleukin 6 (IL-6), to recruit the immune system for clearance and engage this potent physiological anti-tumour response ^5^. Secondly, this approach provides a novel therapeutic opportunity for the targeted killing of senescent cancer cells ^6^. The success of future therapies will rely on identifying targets for senescence induction in cancer cells whose modulation has minimal impact on neighbouring, normal cells, and will likely require tailored therapies on a subtype basis.

The mechanism by which p16 positive BLBC continually evade senescence induction remains elusive. In order to address this question and tackle the current lack of targeted treatment options for this disease subset, we employed a siRNA screening strategy to identify prosenescence drug targets for p16 positive BLBC with the goal of identifying candidate genes with potential clinical utility.

## METHODS

### Cell Culture and Reagents

Unless otherwise stated, all reagents were purchased from Sigma. MDA-MB-468 (ATCC® HTB-132^™^; p16^+/+^, p53^+/+^, p21^+/+^ and RB-null) and MDA-MB-231 cells (ATCC® HTB-26^™^; p16-null, p53^+/+^, p21^+/+^, RB^+/+^) were purchased from ATCC, while HeLa cells were purchased from CRUK. Dulbecco’s Modified Eagles Medium (DMEM) supplemented with 10% foetal bovine serum (FBS, Biosera), 2mM L-Glutamine (Life Technologies) and 1mM Sodium Pyruvate was used for MDA-MB-468 and MDA-MB-231 cells. HeLa cells were maintained in DMEM supplemented with 5% FBS and 2mM L-Glutamine. Normal, finite HMECs were isolated from reduction mammoplasty tissue of a 21-year-old individual, specimen 184, and were cultured in M87A medium as previously described ^7^. Cellular senescence refers to normal, finite HMECs cultured to stasis. Premature senescence refer to cells triggered to enter a premature senescence via siRNA knockdown. Normal human mammary fibroblasts (HMFs) were obtained from reduction mammoplasty tissue of a 16-year-old individual, donor 48 (Stampfer et al. 1981). The cells were seeded at 10,000 cells/cm^2^ and maintained in Dulbecco’s Modified Eagles Medium (DMEM) (Life Technologies, UK) supplemented with 10% foetal bovine serum (FBS) (Labtech.com, UK), 2mM L-glutamine (Life Technologies, UK) and 10μg/mL insulin from bovine pancreas (Sigma). Normal human neonatal foreskin keratinocytes (NFKs) were cultured at 7,500/cm^2^ and maintained in Keratinocyte Serum Free Medium with supplements (Invitrogen). All cells were maintained at 37°C/5% CO_2_. All cells were routinely tested for mycoplasma and shown to be negative.

### Identification of the top six RP hits

Using a data mining approach, we cross referenced phenotypic data from a previously published genome-wide siRNA screen in normal HMECs ^8^ with the results of a similar screen performed in p16 positive HeLa cells (unpublished) to identify genes whose knockdown generated a senescent phenotype only within the HeLa cancer cell context. This exercise identified 57 siRNAs which were selected for further phenotypic screening in HeLa and MDA-MB-468 cells. This list included 2/78 mitochondrial ribosomal proteins (MRPL13 and MRPS24), 8/98 ribosomal proteins (RPL14, RPL18, RPL34, RPL35a, RPLP2, RPS18, RPS3A and RPS7) and UBA52 (ubiquitin A-52 residue ribosomal protein fusion product 1). From this, 25 siRNAs which generated a senescent phenotype in both p16 positive cancer cell models were identified, which included the 11 siRNAs which targeted ribosomal-related proteins. This list included the 11 siRNAs targeting ribosomal-related proteins detailed above.

### siRNA knockdown experiments

Unless otherwise stated all siRNA knockdown experiments were performed as described below and harvested at 5 days-post transfection. The siRNA sequences used are provided in **Supplementary Table 1**. For high content analysis (HCA), cells were reverse transfected with 30nM siRNA pools at a 1:1:1 ratio (Ambion) using HiPerFect (Qiagen) for HeLa, MDA-MB-468s and HMECs or DharmaFect 2 (Dharmacon) for HMFs and NFKs in 384-well format. Reverse transfections were performed at the following passages: HMECs – passage 6 (P6); HMFs – passage 11; and NFKs – P6. Control siRNAs targeting glyceraldehyde-3-phosphate dehydrogenase (GAPDH, Ambion), cyclophilin B (PPIB, siGLO, Dharmacon), polo-like kinase 1 (PLK1, Dharmacon) or Chromobox homolog 7 (CBX7, Ambion) were also included as indicated. Cells were incubated at 37°C/5% CO_2_ and medium changed after 46hr. Cells were then fixed/stained 72hr later and imaged as described below. For immunoblotting or qRT-PCR, cells were seeded in 6-well plate format and harvested for protein or RNA extraction as described below.

### Z score generation

For each of the parameters analysed, significance was defined as three Z scores from the negative control mean. Z scores were generated according to the formula below:

**Z score=**(mean value of two independent experiments for target siRNA **–** mean value (of two independent experiments) for GAPDH siRNA)**/**Standard deviation (SD) for GAPDH siRNA of two independent experiments.

For the phenotypic siRNA screens, each of the data points from two independent experiments were combined and mean values and SDs were calculated. This method generated a highly stringent significance threshold, ensuring only the strongest hits were identified.

### Immunofluorescence microscopy and high content analysis

Cells were fixed with 3.7% paraformaldehyde, permeabilised for 15min using 0.1% Triton X and blocked in 0.25% BSA before 2hr primary antibody incubations at room temperature. Primary antibodies used are listed in **Supplementary Table 2**. Permeabilisation prior to rabbitαNCL incubation was for 30min. Cells were blocked in 1% BSA prior to overnight incubations with rabbitαNCL at room temperature, goatαIL-6 or rabbitαp107 at 4°C. Cells were incubated for 2hr at room temperature with the appropriate AlexaFluor-488 or AlexaFluor-546 conjugated secondary antibody (1:500, Invitrogen), DAPI and CellMask Deep Red (Invitrogen). Images were acquired using the IN Cell 1000 or 2200 automated microscopes (GE) and HCA was performed using the IN Cell Developer software (GE).

### Immunoblotting and Densitometry analysis

Cell were lysed in RIPA buffer supplemented with 4% protease cocktail inhibitor (Roche) and protein concentration was determined using the Bio-Rad Protein Assay kit (Bio-Rad). Lysates were re-suspended in 6X Laemmli Sample Buffer (0.1M Tris pH6.8, 20% glycerol, 1% β-mercaptoethanol, 1% sodium dodecyl sulphate (SDS), 0.01% Bromophenol blue) and used for immunodetection. Primary antibodies used are listed in **Supplementary Table 2**.

Protein separation was achieved by SDS-PAGE on 10-12% polyacrylamide gels and proteins were subsequently transferred to Hybond nitrocellulose membrane (GE) using the Bio-Rad Mini-PROTEAN III system. Membranes were blocked in 5% Milk/PBS-Tween for 1hr before overnight incubation with primary antibody at 4°C with the exception of mouseαp16 which was used at room temperature for 2hr. Following 3X PBS-T washes, membranes were incubated with an appropriate horseradish peroxidase (HRP)-conjugated secondary antibody for 1hr. Bands were then visualised using Enhanced-Chemiluminescence (ECL, GE). Densitometry analysis was conducted using Image J software. Density levels were corrected for protein loading and were expressed relative to the negative siRNA control.

### Quantitative RT-PCR (qRT-PCR)

Primers were purchased from Eurofins Genomics and their sequences are listed in **Supplementary Table 3**. For siRNA knockdown validation, RNA was extracted 76hr post-transfection and gene-expression levels were quantified using target-specific probes. CT values were normalised to endogenous HPRT1 levels and target gene-expression levels were expressed relative to the GAPDH siRNA control for each experiment. All qPCR reactions were run in duplicate for two independent samples.

### qRT-PCR Arrays

Bespoke qRT-PCR arrays to profile the transcripts of 77 ribosomal proteins were purchased from Qiagen. qRT-PCR analysis was performed according to the manufacturer’s instructions and as detailed above. A list of the genes is given in **Supplementary Table S4**.

### Actinomycin D drug treatment

Cells were incubated with 10μg/mL Actinomycin D in DMSO (VWR) for 8hr at 37°C/5% CO_2_ before DAPI/rabbitαNCL immunofluorescence staining and image acquisition using the IN Cell 1000 at 20X magnification.

### Senescence-associated β-galactosidase activity assay

Cells were fixed with 0.2% glutaraldehyde and incubated with 5-bromo-4-chloro-3-indolyl-beta-D-galacto-pyranoside (X-gal) solution for 20hr at 37°C without additional CO_2_. Images were acquired using a light microscope (Nikon) at 20X magnification.

### SYTOX staining assay

Cells were incubated with 500nM SYTOXGreen nucleic acid stain (SYTOX, Invitrogen) and 1.62μM Hoechst 33342 (Invitrogen) for 2.5hr at 37°C/5% CO_2_. Images were acquired using the IN Cell 1000 at 10X magnification.

### Breast Cancer Tissue Microarray Analysis

Tumour and surrounding normal tissue: Tissue samples (snap frozen) were obtained from patients undergoing breast surgery between 2011 and 2013; all patients gave their consent according to the tissue bank protocol. Samples of tumour tissue and surrounding morphologically normal tissue, taken >5cm from the tumour, were obtained from each patient. All samples were obtained from the Barts Cancer Institute Breast Tissue Bank and were covered by Research Tissue Bank Ethics Approval.

### Immunofluorescence staining of breast cancer tissue microarrays

Formalin-fixed, paraffin-embedded primary human BLBC tissue sections (0.5uM) were dewaxed in 100% xylene (2X 3 minute washes) and rehydrated in 100% ETOH (2X 3 minute washes) to distilled water. Antigen retrieval was performed by heating the sections in 10mM citric acid (pH6) at full power (850W) in a domestic microwave oven for 2X 5 minutes. Sections were then permibilised using 0.1% Triton/PBS for 15 minutes before being blocked in 1% BSA/PBS for 1 hour at room temperature. Antibodies were used to detect pan Cytokeratin (AE1/AE3, AB27988, Abcam, 1:1000), p16 (JC8 mouse monoclonal, 1:700), RPS3A (HPA047100, Sigma, 1:1,000) and RPS7 (57637, Abcam, 1:1,000). All antibodies were diluted in 1% BSA/PBS and sections were incubated in primary antibody overnight at 4°C. Sections were then incubated with the appropriate secondary antibodies (AlexaFluor 488 goat anti-mouse or AlexaFluor 546 donkey anti-rabbit, Life Technologies, 1:500) and DAPI (1 μg/ml) diluted in 1% BSA/PBS for 1 hour at room temperature. Sections were mounted under glass coverslips using Immu-mount mounting media (Thermo Scientific) and imaged at 40X magnification using the IN Cell 2200 (GE).

### Statistical analysis

Statistical analysis was performed using GraphPad Prism 6. An unpaired Student’s t-test was used to compare the means of two groups with the exception of day 5 versus day 16 data, where an unpaired Student’s t-test was used. For normalised data, an Ordinary One-way ANOVA followed by a Dunnett’s post-hoc test was used. For breast cancer tissue microarray analysis, the association between RPS3A and RPS7 or p16 protein expression used Spearman’s correlation test. Unless otherwise stated, at least two independent experiments were performed in triplicate. The P-value <0.05 was considered statistically significant.

## RESULTS

### siRNA screening identifies select ribosomal proteins as novel pro-senescence targets for p16 positive cancers

To identify novel pro-senescence therapeutic targets in p16+ BLBC we took advantage of the well-established BLBC *in vitro* model, MDA-MB-468 (p16^+/+^, p53^+/+^, p21^+/+^, RB-null; **Supplementary Figure 1a**) for siRNA screening (see screening schematic Figure 1a). MDA-MB-468 cells were reverse transfected with 30nM siRNA pools targeting 57 shortlisted genes (see Methods Section) alongside siRNAs targeting Glyceraldehyde 3-phosphate dehydrogenase (GAPDH; negative control siRNA) and Chromobox protein homolog 7 (CBX7; positive control siRNA) ^8^. Hit selection was based on two preliminary morphological parameters commonly associated with senescence activation: a cessation of cell proliferation and increased cell area. As hypothesised, CBX7 siRNA triggered a potent senescence-like phenotype in MDA-MB-468 cells (Figure 1b-c, **top panels**). We identified 25 siRNAs that activated a senescence-like phenotype in this model, 11 of which targeted ribosomal proteins (RPs), namely eight ribosomal proteins (RPL14, RPL18, RPL34, RPL35a, RPLP2, RPS18, RPS3A and RPS7), two mitochondrial ribosomal proteins (MRPL13 and MRPS24), and UBA52 (ubiquitin A-52 residue ribosomal protein fusion product 1). Kaplan Meier analysis highlighted the poor outcome for BLBC associated with elevated p16 expression, and revealed that elevated expression of six of the RPs (RPL14, RPL18, RPL34, RPL35A, RPS3A and RPS7) is predictive of a poor prognosis in BLBC (Figure 1d). No significant impact on prognosis for BLBC was observed for RB, p107 or p130 (not shown). By contrast, similar analysis for 10 RPs whose knockdown did not impact cancer cell proliferation (see Methods) revealed that elevated expression had no significant impact on BLBC prognosis or, in one case (RPS23), was protective (**Supplemental Figure 1b**). This suggests that the 6 RP hits may have a predictive power for BLBC. Therefore, we prioritised these six, disease relevant RPs for multiparameter phenotypic analysis which revealed the induction of a robust senescent-like phenotype in MDA-MB-468 cells (Figure 1b-c, **lower panels**). Next, we further validated the potential merit of RP knockdown for senescence induction in HeLa cells (p16+ cervical cancer cell line, lacking function p53 and RB), demonstrating the activation of a senescence-like phenotype (Figure 1e, **Supplementary Figure 1c**). Following validation of mRNA knockdown for the six RP siRNA pools in MDA-MB-468 cells (**Supplementary Figure 1d**), we deconvoluted each siRNA pool performing multiparameter phenotypic validation to control for off-target effects. This identified three functional siRNAs for RPL14, RPL18, RPL35A and RPS3A, two for RPS7, and a single RPL34 siRNA that generated a robust senescence response (Figure 1f and data not shown). This was subsequently confirmed at the protein level for individual siRNAs targeting RPS3A and RPS7 (**Supplementary Figure 1e, see also later** Figures 5).

**Figure 1.**
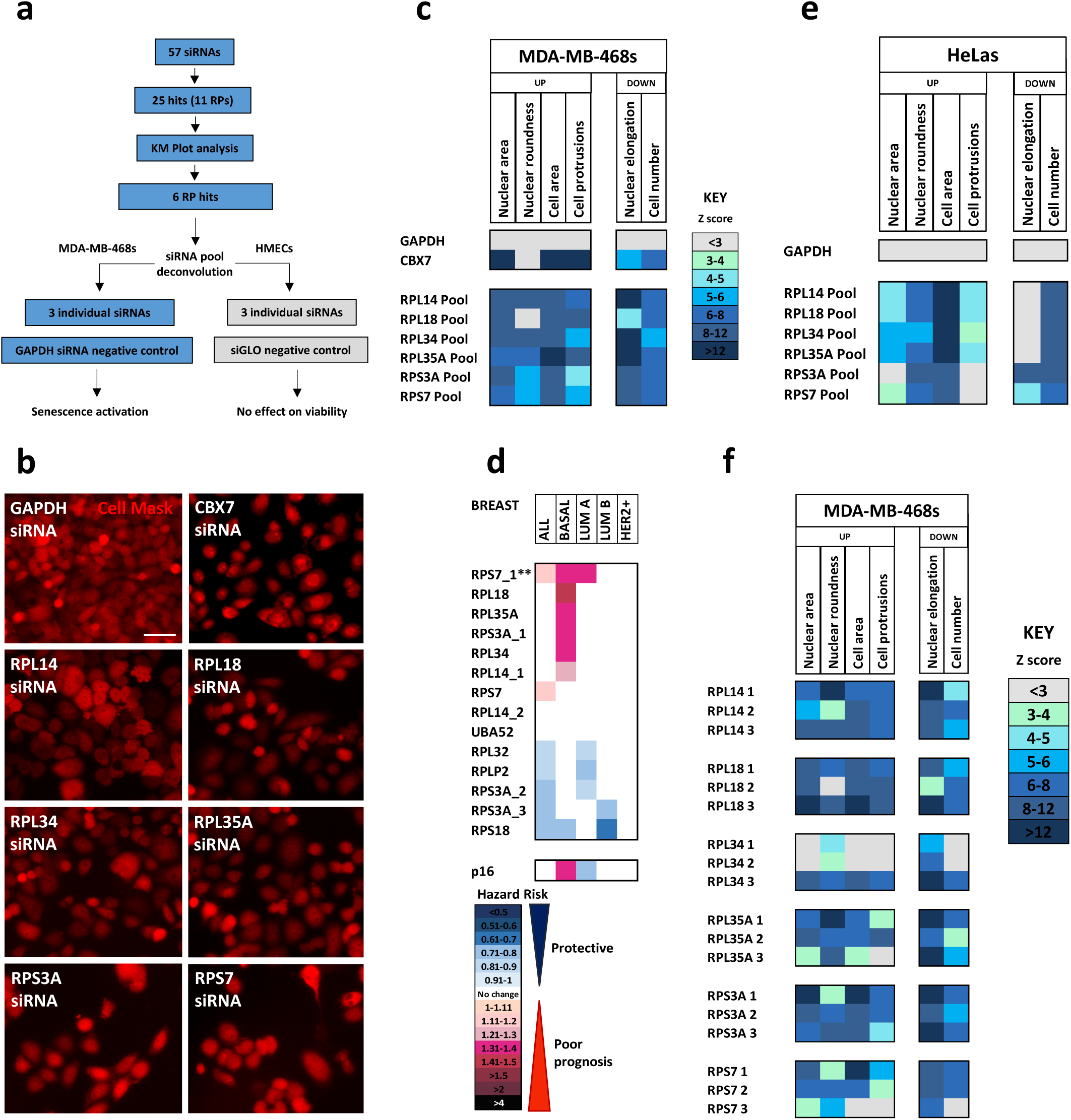
Pro-senescence siRNA screening: knockdown of six ribosomal proteins drives senescence in p16-positive cancer cells. **(a)** Schematic depicting the siRNA screening strategy conducted in MDA-MB-468 cells and HMECs. **(b)** Representative immunofluorescence images of MDA-MB-468 cells transfected with GAPDH siRNA, CBX7 siRNA or each of the RP siRNA pools prioritised for further validation (Cell Mask, red). Size bar 50µm. **(c)** Heatmap depicting mean Z-scores for six senescence-associated morphological parameters following RP siRNA knockdown and multiparameter analysis in MDA-MB-468 cells**. (d)** Heatmap displaying the Hazard Risk for 10 year relapse free survival with a logrank p value <0.05 for ten RPs and p16 for All breast cancer cases (N=3,955), Basal-like (N=580), Luminal A (N=1,764), Luminal B (N=1,002) and HER2+ (N=335). Data generated from KMPLOT for median expression with the exception of RPS7_1 where lower tertile versus upper tertiles is shown (**). **(e)** As stated in (c) performed in HeLa cells. **(f)** Heatmap depicting Z-scores for six phenotypic validation of the six RP siRNA pools following siRNA pool deconvolution in MDA-MB-468 cells.

An important feature of future pro-senescence therapies is that they should not affect the viability nor trigger a senescence response in neighbouring, normal cells. Thus, normal, non-immortalised, finite lifespan human mammary epithelial cells (HMECs) ^7^ were transfected with the original siRNA pools or three individual siRNAs targeting each of the six RPs. Unlike the cancer cell setting, RP knockdown did not alter proliferation or activate a senescence programme in normal HMECs (**Supplementary Figure 1f**), indicating that the induction of a senescence-like programme following specific ribosomal targeting may be a cancer cell-specific response. To further test this hypothesis, we performed phenotypic screening in HMFs (a different breast tissue cell type) as well as NFKs (epithelial cells from a different tissue). In line with the findings within HMECs, RP knockdown did not alter the proliferation or activate a senescence programme within these disparate, normal cell types (**Supplementary Figure 1g-h**).

### RP knockdown results in activation of a panel of senescence markers, altered nucleolar morphology and a stable cell cycle arrest

Next, we sought to fully characterise the consequence of senescence initiation within the cancer cell context using a panel of well-established senescence markers. This revealed that RP knockdown resulted in a significant decrease in the percentage of Ki67 positive cells (proliferation marker, **Supplementary Figure 2a-b**), a significant increase in the SASP factors at both the transcript level (IL-1α, IL-1β and IL-6 Supplementary figure 2c-e) and at the protein level for IL-6 (Figure 2a, RPL14 p=0.059) and IL-8 (Figure 2b, RPL18, RPS3A and RPS7), and an increase in SA-β-gal activity (**Supplementary Figure 2h**). Importantly, knockdown of the RPs did not result in an altered DNA damage response as assessed by 53BP1 foci multiparameter analysis (Figure 2c-d). Nucleoli, the site of ribosomal biogenesis, are commonly enlarged and irregular in tumour cells, and this is associated with a poor prognosis in breast cancer ^9^. In addition, nucleolar dysfunction following Nucleolin (NCL) knockdown in HeLa cells, primary human fibroblasts ^10^ or HMECs ^8^, generates a single, enlarged nucleolus and induces senescence. Therefore, we sought to examine the impact of RP knockdown on nucleolar abundance and morphology, with a view to prioritising only those RP siRNAs which did not influence this nuclear feature in normal HMECs. Initially, the nucleolar signature for HMEC cellular or premature senescence (following transfection with 30nM CBX7 siRNA) was established using HCA (**Supplementary Figure 2i-j, respectively**). p16 positive cellular senescent normal HMECs displayed a single enlarged nucleolus, as observed previously ^10^. Intriguingly, induction of the p16 positive premature senescence programme resulted in a highly irregular nucleolar signature, associated with a significant increase in nucleoli number, area and elongation (Figure 2e), which was distinguishable from nucleolar disassociation following treatment with actinomycin D (**Supplementary Figure 2k**). Together, this established the nucleolar profile of cellular senescent (single enlarged nucleoli) and prematurely senescent (enlarged, irregular and elongated nucleoli) HMECs and identified a marker capable of distinguishing between these two differing routes to senescence activation. Importantly, RP knockdown in HMECs using either RP pools (Figure 2e) or individual RP siRNAs (data not shown) did not result in any significant alterations to nucleoli number or morphology, suggesting that specific RP silencing in these cells does not induce significant nucleolar dysfunction, and provided further evidence that knockdown of our six RPs is well-tolerated in normal HMECs. By contrast, both CBX7 and RP knockdown in MDA-MD-468s generated a marked change in nucleolar phenotype (increase in nucleolar abundance and morphology), reminiscent of the premature senescence signature (Figure 2f-g).

**Figure 2.**
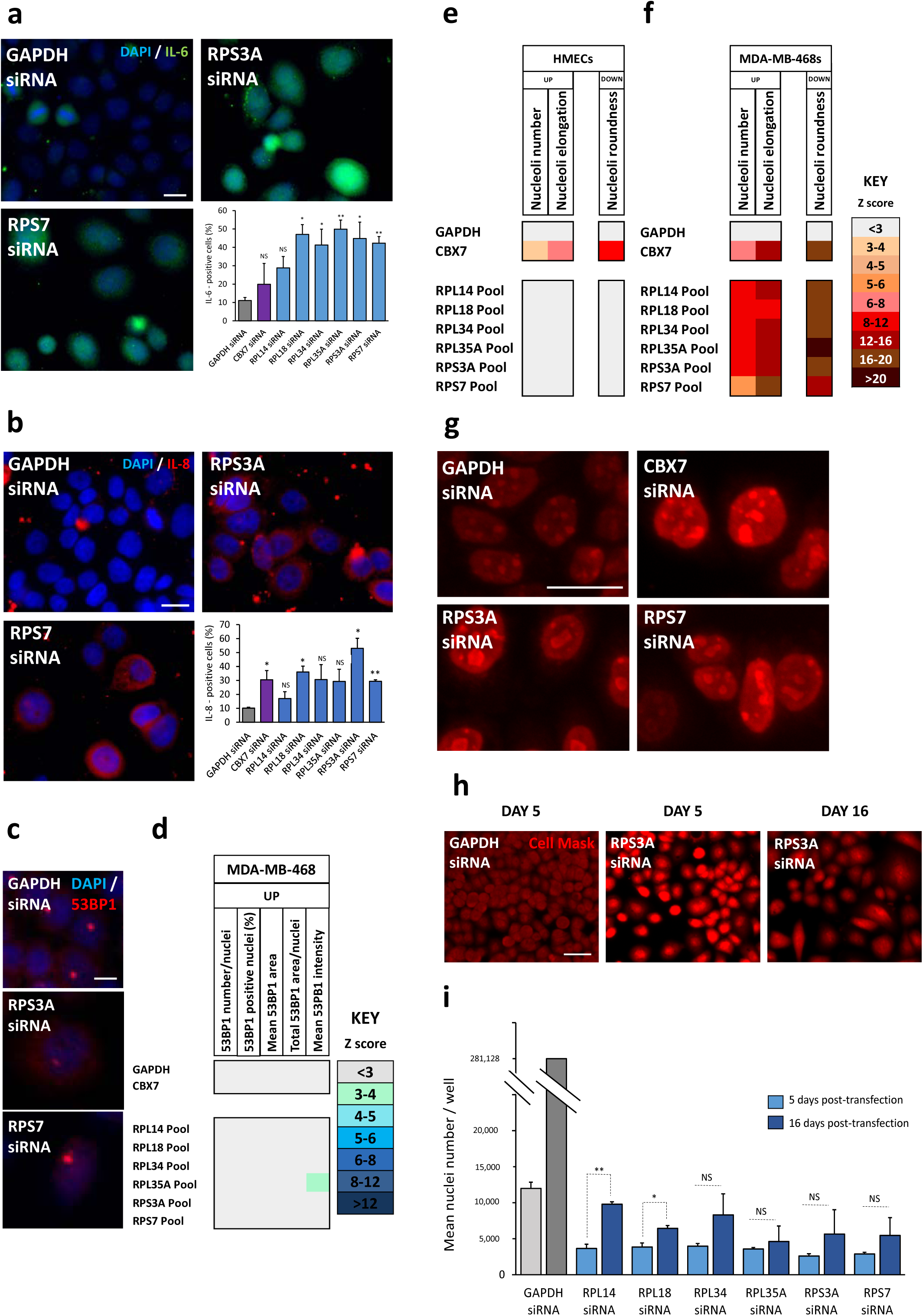
RSIS is stable and is characterised by an altered nucleolar morphology and activation of the senescent-associated secretory phenotype (SASP). **(a)** Representative immunofluorescence images of MDA-MB-468 cells transfected with GAPDH or 3nM siRNA targeting RPS3A or RPS7 (DAPI, blue; IL-6, green). Size bar 50µm. Bar chart showing the mean +SD percentage of IL-6-positive cells. **(b)** Heatmap depicting the mean Z score for each of the 53BP1 parameters selected for quantitation following transfection of MDA-MB-468 cells with siRNA targeting GAPDH, CBX7 or each of the six RPs. **(c-d)** Heatmap depicting the mean Z score for each of the nucleolar parameters selected for quantitation in HMEC cells (c) and MDA-MB-468 cells (d). **(e)** Representative immunofluorescence images of MDA-MB-468 cells transfected with GAPDH, CBX7, RPS3A or RPS7 siRNA (Nucleolin (NCL), red). Size bar 25µm. **(f)** Representative immunofluorescence images of MDA-MB-468 cells transfected with GAPDH or RPS3A siRNA at 5 or 16 days post-transfection (Cell Mask, red; N=1). Size bar 50µm. **(g)** Bar chart showing mean +SD nuclei number at 5 or 16 days post-transfection following three independent experiments each performed in triplicate. Cell numbers in GAPDH siRNA treated wells could not be quantified as the wells were overgrown. Instead, day 16 cell number for the GAPDH siRNA control was calculated based on known population doubling times. *=p<0.05, **=p<0.01, ***=p<0.001, ****=p<0.0001.

Another key feature of the senescence program is a stable cell cycle arrest. Therefore, the long-term stability of the RP knockdown phenotype was examined in MDA-MB-468 cells (Figure 2h-i). Knockdown of RPL14 or RPL18 did not result in a completely stable cell cycle arrest and was associated with a modest but significant (p<0.05) increase in cell number over a 16 day time course when compared to the GAPDH negative control. However, knockdown of RPL34, RPL35A, RPS3A or RPS7 resulted in a highly stable cell cycle arrest. In summary, we have identified and validated six RPs whose knockdown engages the senescent programme in p16 positive cancer cell models whilst being well tolerated in a panel of normal human cell types (HMECs, HMFs and NFKs). The initiation and maintenance of senescence is defined by a panel of well-established senescence markers, including the SASP factors IL-6 and IL-8, combined with a nucleolar signature which aligns with premature rather than cellular senescence. We define this trigger to cancer cell-specific senescence as ribosomal stress-induced senescence (RSIS).

### Ribosomal stress-induced senescence (RSIS) initiation requires endogenous p16 and results in p16 nuclear translocation and p107 upregulation

Next, we sought to explore the mechanism of RSIS initiation, and first determined the status of a key senescence effector, p53, and its downstream target, p21 ^11^. RP knockdown did not significantly alter p53 protein level (MDA-MB-468 cells, p53 positive, **Supplementary Figure 3a**; and pRb-negative) or nuclear abundance (data not shown), and resulted in a decrease in p21 protein levels for each RP pool as assessed by HCA (data not shown) and by western blot following RPS3A and RPS7 knockdown (**Supplementary Figure 3b**). Furthermore, RSIS induction in HeLa cells (p53-negative) was not associated with the activation or stabilisation of p53 (data not shown), further indicating that RP knockdown in a p16 positive context drives senescence induction independently of the p53 axis. Given this, we hypothesised that in p16 positive cancer cells RSIS may be established via re-sensitisation to the endogenously and constitutively expressed p16 protein independent of RB. Whilst total p16 protein levels remained stable following RSIS initiation (Figure 3a), there was a significant increase in the nuclear to cytoplasmic ratio of p16 following knockdown of each of the six RPs, suggesting increased translocation of p16 from the cytoplasm to the nucleus, the site for CDK4/6-Cyclin D complex inhibition (Figure 3b-c, **Supplementary Figure 3c**). Given that the inactivation of RB family members confers a growth advantage to multiple types of tumours, we next determined the consequence of RP knockdown for the CDK/cyclin targets retinoblastoma-like 1 (RBL1/p107), and retinoblastoma-like 2 (RBL2/p130). Knockdown of RPS3A and RPS7 resulted in a significant increase in total p107 and p130 protein (**Supplementary Figure 3d**). Further, knockdown of each of the RP hits generated a significant increase the percentage of p107-postive nuclei (Figure 3d-e, **Supplementary Figure 3e**) and an increase in nuclear levels of p107 within the positive population when compared to the GAPDH negative control (Figure 3f, **Supplementary Figure 3f**). To further test the role of the p16/p107-p130 axis in RSIS initiation, we performed p16 rescue experiments in MDA-MB-468 cells. In each instance, p16 knockdown rescued cell proliferation as well as the morphological alterations (increase in cell and nuclear area) associated with the RSIS programme (Figure 3g-h, **Supplementary Figure 3g**), indicating that RSIS initiation is dependent on endogenous p16. In summary, we find that RSIS maintenance in p16 positive cancer models occurs via a p53/p21-independent mechanism, and that endogenously expressed p16 is necessary and sufficient to activate the p16/p107-p130 axis to initiate and maintain the RSIS programme.

**Figure 3.**
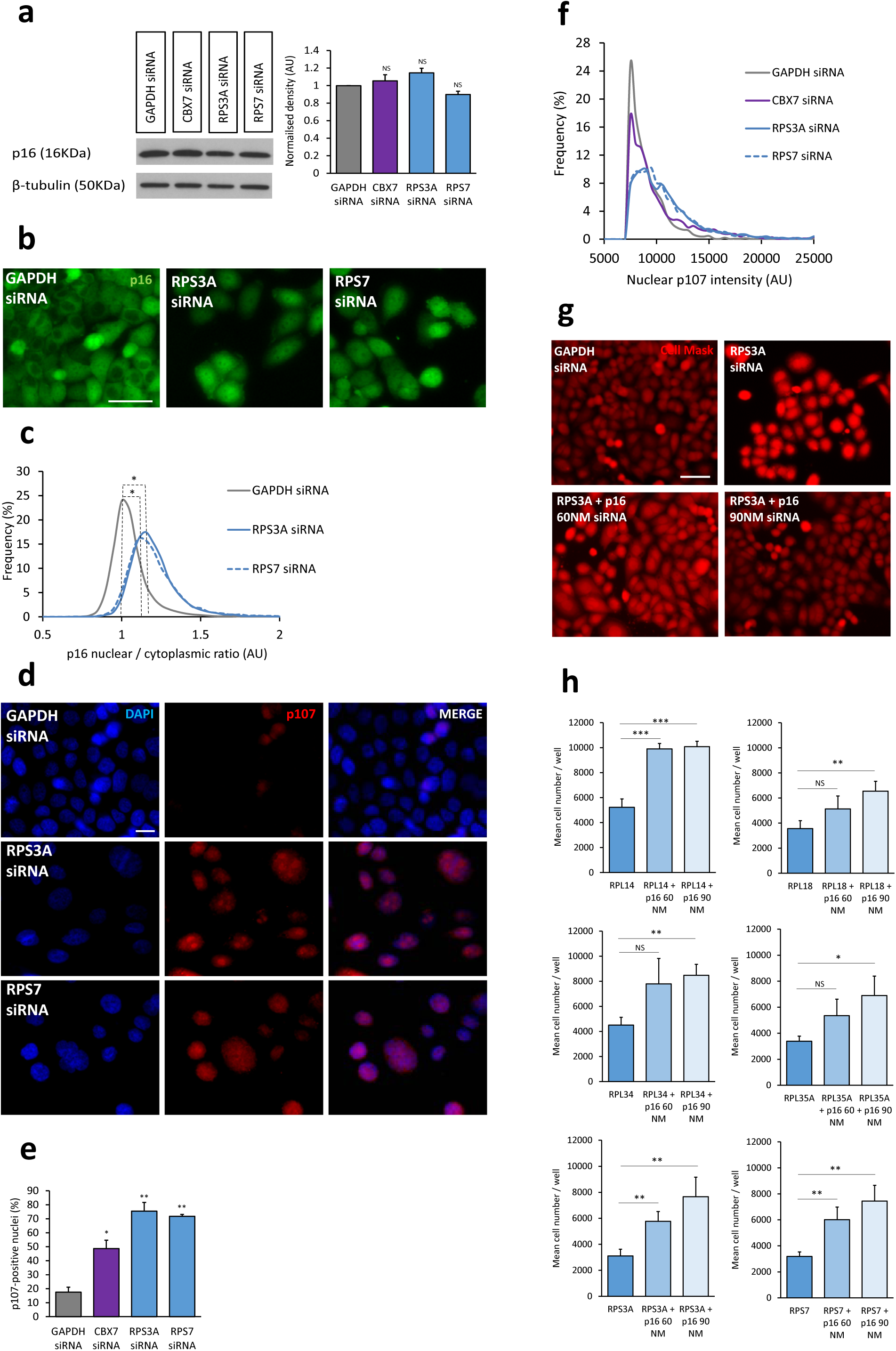
RSIS initiation requires endogenous p16 and results in p16 nuclear translocation and p107 upregulation. **(a)** Western blot analysis of p16 in MDA-MB-468 cells following transfection with GAPDH, CBX7, RPS3A or RPS7 siRNA. Loading control: β-tubulin (left). Densitometry analysis of p16 (right). Bars denote mean density levels +SD normalised to GAPDH siRNA. **(b)** Representative immunofluorescence images of MDA-MB-468 cells transfected with GAPDH, RPS3A or RPS7 siRNA (p16, green). Size bar 50µm. **(c)** Representative frequency distribution depicting the p16 nuclear/cytoplasmic ratio at 5 days post-transfection following GAPDH, RPS3A or RPS7 siRNA knockdown in MDA-MB-468 cells. **(d)** Representative immunofluorescence images of MDA-MB-468 cells transfected with GAPDH, RPS3A or RPS7 siRNA (DAPI, blue; p107, red). Size bar 50µm. **(e)** Bar chart depicting mean +SD percentage of p107-positive nuclei. **(f)** Representative frequency distribution depicting p107 intensity levels within the positive population following GAPDH, CBX7, RPS3A or RPS7 siRNA knockdown in MDA-MB-468 cells. **(g)** Representative immunofluorescence images of MDA-MB-468 cells transfected with 90nM GAPDH, 3nM RPS3A alone or 3nM RPS3A with 60nM or 90nM p16 siRNA (Cell Mask, red). Size bar 50µm. **(h)** Bar charts depicting mean +SD cell number following RP knockdown or rescue with p16 siRNA 5 days post transfection. Three independent experiments were performed, each containing three technical replicates. *=p<0.05, **=p<0.01, ***=p<0.001, ****=p<0.0001.

### In the absence of p16, RP knockdown results in p53 stabilisation and Caspase-3 mediated apoptosis

To further explore the role of p16 in RSIS initiation, a second BLBC model, MDA-MB-231 (p16 negative, p53 positive, p21 positive, **Supplementary Figure 1a**) was selected for RP knockdown. Initially, CBX7 siRNA was used to assess whether senescence could be triggered in this model. Despite their p16 status, CBX7 knockdown in MDA-MB-231 cells triggered a senescence-like phenotype, characterised by a significant decrease in proliferation together with an increase in cell and nuclear area (**Supplementary Figure 4a-b**). In addition, arrested MDA-MB-231 cells were associated with a nuclear stabilisation of p21 (**Supplementary Figure 4c**), and p21 siRNA rescue experiments showed that senescence initiation was dependent on endogenous p21 (**Supplementary Figures 4d**). Together, these findings suggest that in the absence of p16, CBX7 knockdown re-sensitises MDA-MB-231 cells to endogenous p21, triggering a p21-dependent senescence programme.

Subsequently, RP knockdown was performed in the MDA-MB-231 model. Surprisingly, and in contrast to CBX7 knockdown, initial observations indicated that RP knockdown activates a death-like phenotype when compared to the GAPDH negative control (Figure 4a, **Supplementary Figure 4e**), characterised by a significant reduction in cell number and an increase in nuclear roundness for each of the six RP hits (data not shown). Next, we showed that RP knockdown was accompanied by a significant increase in the number of SYTOX positive nuclei (nucleic acid binding dye that penetrates apoptotic cell membranes), confirming cell death activation in this model (Figure 4b, **Supplementary Figure 4f**). We further probed the mechanism by which RP knockdown in a p16 negative context induced cell death and found that RPS3A and RPS7 knockdown induced a significant increase in p53 protein (Figure 4c-d). Previous work has demonstrated a role for a 14kDa cleavage product of p21 in p53-mediated apoptosis ^12^, however, despite repeated attempts we were unable to detect this C-terminal cleavage product using an N-terminal specific p21 antibody (data not shown). However, RPS3A and RPS7 knockdown resulted in an increase in pro-caspase 9, and cleaved caspase 9 (Figure 4e), with knockdown of each of the six RPs resulting in a significant increase in the percentage of caspase-3 positive nuclei (p53-mediated apoptosis markers; Figure 4e, **Supplementary Figure 4g**). Predictably, CBX7 siRNA did not induce any significant changes to SYTOX or caspase-3 staining and accords with our previous data suggesting that following CBX7 knockdown, MDA-MB-231 cells enter a p21-mediated senescence programme. Taken together, we find a divergence in response to CBX7 versus RP knockdown in a p16 negative context and demonstrate that RP knockdown drives apoptosis activation via a p53-caspase 9/3 mediated mechanism in the absence of p16.

**Figure 4.**
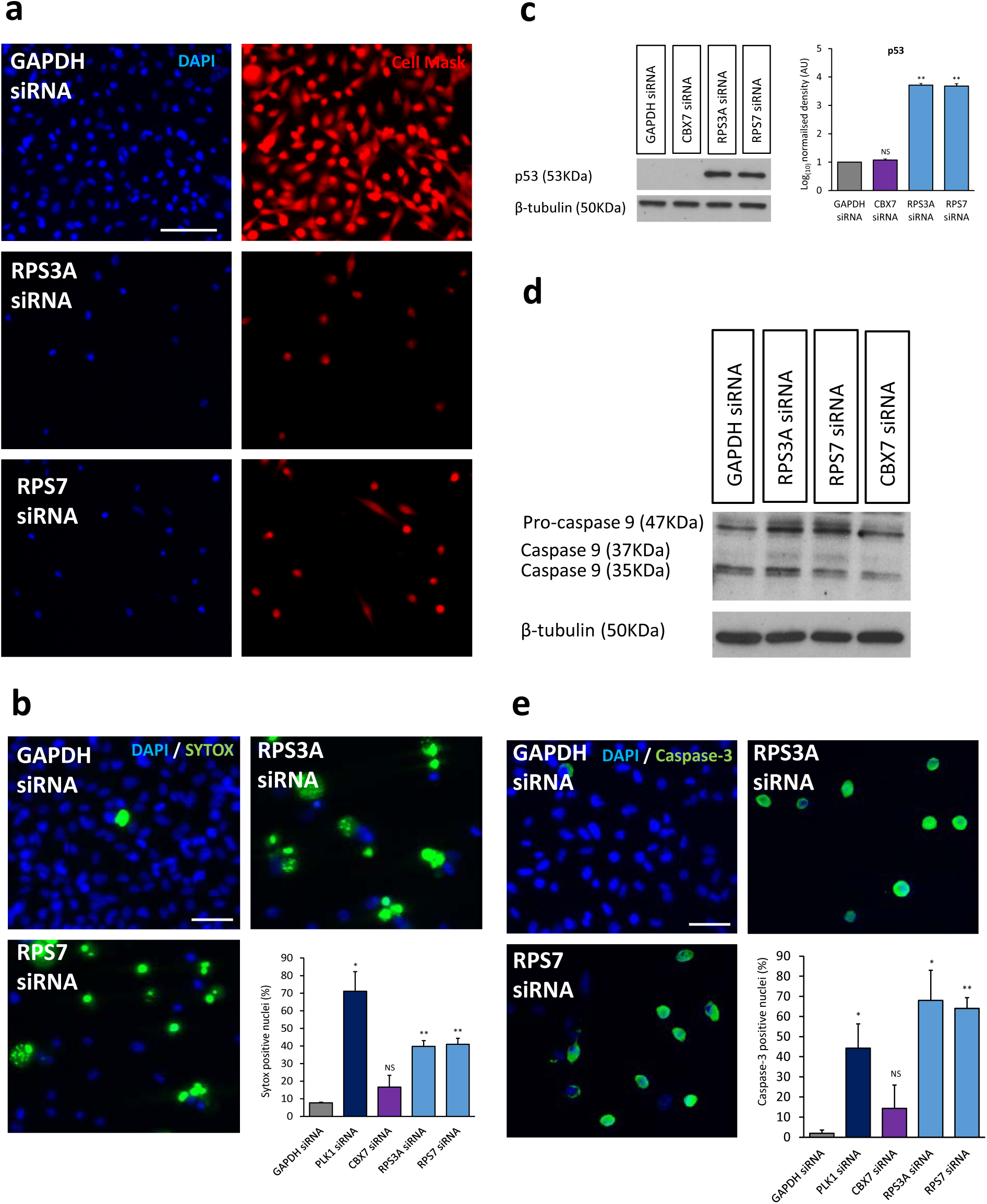
In the absence of p16, RP knockdown results in p53 stabilisation and Caspase-3 mediated apoptosis. **(a)** Representative immunofluorescence images of MDA-MB-231 cells transfected with GAPDH, RPS3A or RPS7 siRNA (DAPI, blue; Cell Mask red). Size bar 50µm. **(b)** Representative immunofluorescence images of MDA-MB-231 cells transfected with GAPDH, RPS3A or RPS7 siRNA (Hoescht 33342, blue; SYTOX green). Cells were stained at 6 days post-transfection and live cell imaging performed. Size bar 50µm. Bar chart depicting mean +SD percentage of SYTOX-positive cells. **(c)** Western blot analysis of p53 in MDA-MB-231 cells. Cells were reverse transfected with GAPDH, CBX7, RPS3A or RPS7 siRNA and protein was harvested at 5 days post-transfection. Loading control: β-tubulin (left). Densitometry analysis of p53 (right). Bars denote mean Log_(10)_ density levels +SD normalised to GAPDH siRNA. **(d)** Western blot analysis of Pro-caspase 9 and cleaved Caspase 9 in MDA-MB-231 cells. Cells were reverse transfected with GAPDH, CBX7, RPS3A or RPS7 siRNA and protein was harvested at 5 days post-transfection. Loading control: β-tubulin, N=1. **(e)** Representative immunofluorescence images of MDA-MB-231 cells transfected with GAPDH, RPS3A or RPS7 siRNA (DAPI, blue; Caspase-3, green). Size bar 50µm. Bar chart depicting mean +SD percentage of Caspase-3-positive cells. *=p<0.05, **=p<0.01, ***=p<0.001, ****=p<0.0001.

### Reciprocal relationship between RPS3A and RPS7 during RSIS and in BLBC

RPS3A and RPS7 knockdown consistently produced the most potent RSIS phenotype as assessed by multiple measures and, therefore, we prioritised these two RPs to explore the impact of RSIS on 77 ribosomal transcripts in MDA-MB-468 cells using qPCR arrays (see **Supplementary Table 4**). This revealed that both routes to RSIS generated a common RP expression signature (Figure 5a). Whilst the majority of ribosomal transcripts (69/77) did not significantly alter following RPS3A or RPS7 knockdown, intriguingly, knockdown of RPS3A resulted in a significant reduction in *RPS7* and *RPL14* transcript levels. Similarly, knockdown of RPS7 resulted in a significant suppression of *RPS3A* and *RPL14*. Finally, both RPS3A and RPS7 knockdown resulted in a significant upregulation of the same 4/77 RPs, namely *RPL10A, RPL17, RPS2 and RPS6KA1*. These results show that RPS3A or RPS7 knockdown does not lead to a global suppression of RP expression, but that RSIS uncouples RP transcript homeostasis driving the dysregulation of a subset of transcripts.

**Figure 5.**
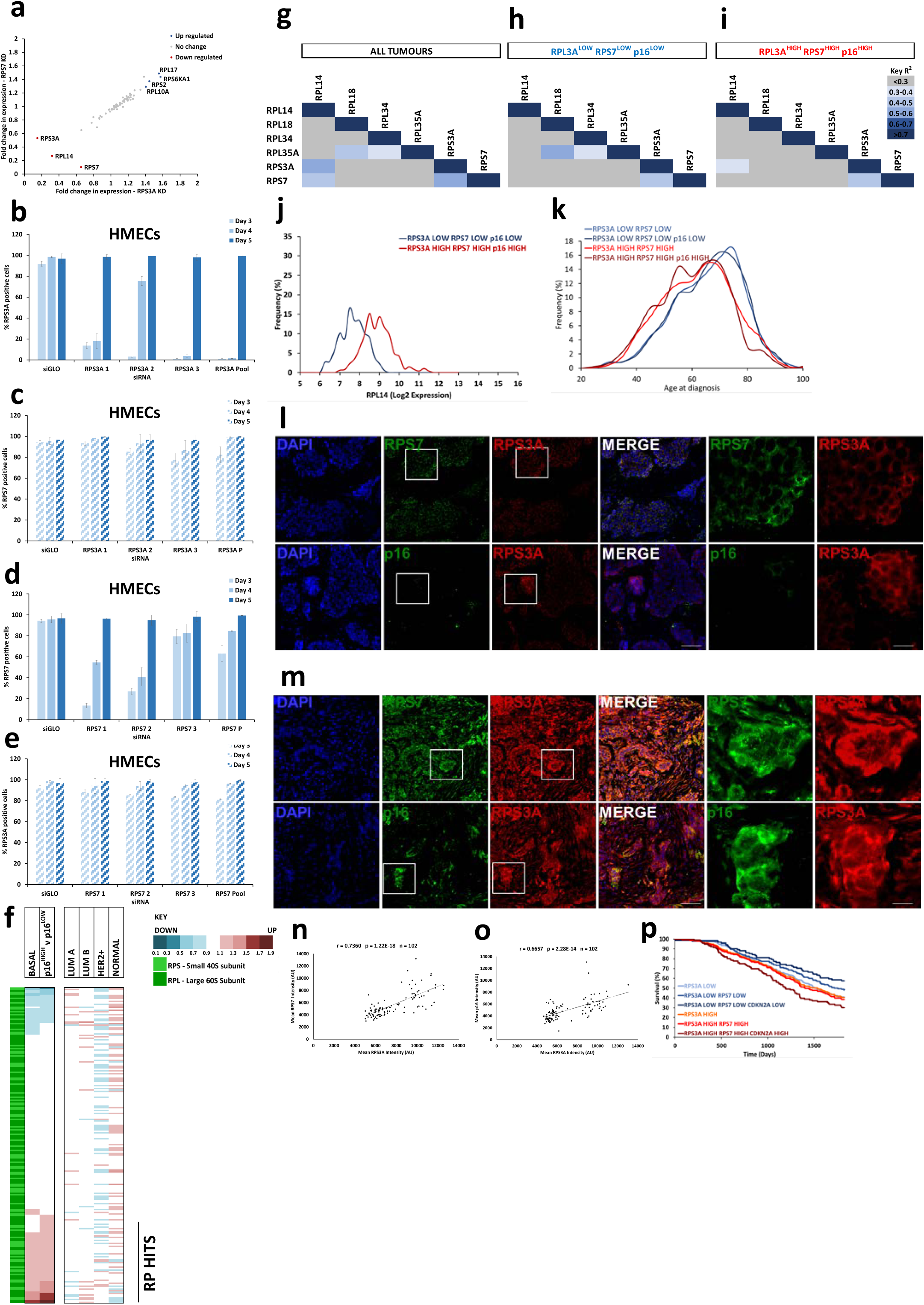
Expression of ribosomal proteins in selectively dysregulated in breast cancer. Elevated expression of the six ribosomal proteins is associated with a poor prognosis in basal-like breast cancer, and a RPS3A^HIGH^RPS7^HIGH^p16^HIGH^ signature results in a particularly poor outcome for patients with breast cancer. **(a)** Log2 Fold change in expression of 77 ribosomal transcripts following transfection of MDA-MB-468 cells with RPS3A or RPS7 siRNA and profiled by qRT-PCR Array at 72hr post transfection. Significantly up regulated (blue) and down regulated (red) transcripts are indicated. **(b-e)** HMECs were transfected with RPS3A 1,2,3 or pool (b-c) or RPS7 1,2,3 or poo (d-e). Cells were fixed at Day 3, 4 and 5 post transfection, and high content image analysis performed for RPS3A and RPS7. Bar charts showing % (b-c) percentage RPS3A and RPS7 positive cells following RPS3A knockdown and (d-e) percentage RPS7 and RPS3A positive cells following RPS7 knockdown are shown. **(f)** Heatmap depicting up (red) or down (blue) regulation of ribosomal proteins (RPS, light green; RPL, dark green) for the PAM50 subtypes (Basal, N=329; Luminal A, N=718; Luminal B, N=488; Her2+, N=240; and Normal N=199) and for p16^HIGH^ tumours irrespective of PAM50 subtype. **(g-i)** Heatmap for R^2^ values for Log_2_-fold change gene expression values each of the six RPs in (g) all tumour samples (h) RPS3A^LOW^RPS7^LOW^p16^LOW^ tumours and (i) RPS3A^HIGH^RPS7^HIGH^p16^HIGH^ tumours, all irrespective of PAM50 subtype. **(j)** Frequency distribution Log_2_-fold expression for RPL14 expression in RPS3A^LOW^RPS7^LOW^p16^LOW^ (blue) and RPS3A^HIGH^RPS7^HIGH^p16^HIGH^ (red) tumours. **(k)** Frequency distribution of age at diagnosis for RPS3A^LOW^RPS7^LOW^ (light blue), RPS3A^LOW^RPS7^LOW^p16^LOW^ (dark blue), RPS3A^HIGH^RPS7^HIGH^ (light red), and RPS3A^HIGH^RPS7^HIGH^p16^HIGH^ tumours (dark red). **(l-o)** Tissue microarrays containing 102 breast cancer tissue samples were stained with DAPI (blue) and either RPS7 (green)/ RPS3A (red) or p16 (green) / RPS3A (red), and high content image analysis performed to determine the average intensity for each of these markers on a per tumour basis. Representative images for a (l) RPS3A^LOW^RPS7^LOW^p16^LOW^ and (m) RPS3A^HIGH^RPS7^HIGH^p16^HIGH^ are shown. (n-o) Scatter plots for the average intensities for each tumour are shown for (n) RPS3A versus RPS7 and (o) RPS3A versus RPS7. The results of Spearman’s rank correlation analysis was also shown. **(p)** Kaplan Meier plot for 5 year risk of death for disease from RP3A^LOW^ (light blue), RPL3A^HIGH^ (orange), RPS3A^LOW^RPS7^LOW^ (midblue), RPS3A^HIGH^RPS7^HIGH^ (red), RPS3A^LOW^RPS7^LOW^p16^LOW^ (dark blue) and RPS3A^HIGH^RPS7^HIGH^p16^HIGH^ (dark red).

We further explored the consequences of RPS3A and RSP7 knockdown at the protein level following RSIS. We find that RPS3A knockdown resulted in a reciprocal reduction in RPS7 protein abundance and vice versa in both MDA-MB-468s (**Supplementary Figure 5a-d**) and MDA-MB-231 (**Supplementary Figure 5e-f**), suggesting that the expression of RPS3A and RPS7 are intimately linked at both the transcript and protein level. To probe this further, we investigated the kinetics of the RPS3A and RPS7 protein levels by performing a timecourse in normal HMECs. Despite potent protein knockdown of RPS3A using individual or pooled siRNAs (Figure 5b), this did not generate a reciprocal reduction in RPS7 in HMECs (Figure 5c), nor was a reduction in RPS3A observed following RPS7 knockdown (Figure 5d-e). Thus, we find that in the absence of RSIS the RPS3A/RPS7 reciprocal relationship is not maintained. This divergence between cancer cells and normal cells led us to explore the relationship between each RP transcript within breast cancer to determine if a similar relationship was present. We determined the expression levels of each RP transcript in the METABRIC dataset in BLBC and in p16^HIGH^ tumours, irrespective of the classical PAM50 subtype (Figure 5f). This revealed coordinated dysregulation of a similar subset of ribosomal proteins in BLBC and p16^HIGH^ breast cancer. Importantly, the RP hits identified within the original siRNA screen (including RSP3A and RPS7) were shown to be co-ordinately upregulated in both BLBC and p16^HIGH^ tumours. Interestingly, each of the other PAM50 subtypes exhibited a unique, subtype dependent RP dysregulation signature.

### Elevated expression of the six ribosomal proteins is associated with a poor prognosis in basal-like breast cancer, and a RPS3AHIGH-RPS7HIGH-p16HIGH signature results in a particularly poor outcome for patients with breast cancer

Next, we sought to determine the consequences for the dysregulation of our RP hits in other cancers. We extended our original Kaplan Meier analysis (Figure 1d, plots for RPS3A and RPS7 are shown in **Supplemental Figure 5g**) to lung and gastric cancers, in which elevated p16 expression is also associated with a poor outcome, together with ovarian cancer (**Supplementary Figure 5h**). Elevated expression of each of our original RP hits resulted in a poor outcome for lung and gastric cancer, in particular for individuals with lung cancer who had never smoked. Strikingly, there was virtually no consequence for altered RP expression in ovarian cancer, nor does p16 appear to influence outcome by any sub-classification applied. Taken together, this data suggested that in the cancer cell setting RPS3A and RPS7 protein abundance may be intimately linked, and prompted us to determine the relationship between our six RPs within breast cancer, irrespective of PAM50 subtype. This revealed a strong correlation (R^2^=0.5668) for RPS3A and RPS7 mRNA expression within breast cancer (**Supplementary Figure 5i**), and cross comparison between each of the six RPs revealed that RPS3A and RPS7 had the strongest relationship (Figure 5g). Given that RPS3A and RPS7 emerged as the most potent hits from our phenotypic screen, combined with the strength of the correlation within the METABRIC dataset, we extended this analysis for RPS3A^LOW^RPS7^LOW^p16^LOW^ (triple LOW) and RPS3A^HIGH^RPS7^HIGH^p16^HIGH^ (triple HIGH) tumours, again irrespective of subtype. Interestingly, we find that our six RPs hits diverge into two distinct groups: in the triple LOW tumours the expression of RPL18, RPL34 and RPL35A correlate, whilst in the triple HIGH tumours RPS3A and RPS7 are linked to RPL14 (Figure 5h-i), the later echoing the relationship between RPS3A, RPS7 and RPL14 observed following RSIS (Figure 5a). Indeed, there is a marked elevation of RPL14 expression in the triple HIGH tumours compared to the triple LOW (Figure 5j) which is not observed for RPL18, RPL34 or RPL35A (**Supplementary Figure 5j-l**). Of relevance, we find that both the RPS3A^HIGH^RPS7^HIGH^ as well as the triple HIGH signature results in an earlier age at diagnosis (Figure 5k), suggesting that even in those tumours which are not p16 positive the co-ordinated dysregulation of RPS3A and RPS7 may drive an earlier onset of disease.

Next, we extended our analysis, determining the R^2^ correlation values for the six RPS and each member of the ribosome in either the triple LOW or triple HIGH tumours (**Supplemental Figure 5m**). This analysis generated three clusters reminiscent of the divergence between RPS3A-RPS7-RPL14 (Cluster 1) and RPL18-RPL34-RPL35A (Cluster 2), together with a considerable number of ribosomal genes which showed no correlation between any of the six RP hits (Cluster 3). Examples of the expression levels for the Cluster 1 member, RPL15, and the Cluster 3 member, RPL10, for triple LOW and triple HIGH tumours is provided (**Supplementary Figure 5n-o**).

Finally, the clinical relevance of the correlation between RPS3A, RPS7 and p16 transcript was examined at the protein level using a tumour tissue microarray containing 102 clinical breast tumours. Significant, positive correlations were found between RPS3A and RPS7 (r = 0.7360; p = 1.22E-18) and between RPS3A and p16 (r = 0.6657; p = 2.28E-14) (Figure 5l-o). The Kaplan Meier plot presented in Figure 5p illustrates that the RPS3A-RPS7 signature collaborates with p16 such that RPS3A^LOW^RPS7^LOW^p16^LOW^ is additively protective whilst RPS3A^HIGH^RPS7^HIGH^p16^HIGH^ tumours have the poorest predicted outcome. In summary, we have identified RSIS as a potential novel route for future pro-senescence therapy. Our top hits, RPS3A and RPS7, have a reciprocal relationship in cancer cells *in vitro*, and this is mirrored at both the transcript and protein level within clinical breast tumour samples. We discover that, irrespective of subtype, RPS3A, RPS7 and p16 are co-ordinately dysregulated, and this signature is associated with a younger age at diagnosis, and that RPS3A and RPS7 synergises with a p16^HIGH^ signature to drive a particularly poor outcome.

## DISCUSSION

Using phenotypic screening, we identified six novel pro-senescence targets whose knockdown triggered RSIS, re-sensitising cancer cells to endogenously expressed p16 to engage a conserved checkpoint. Features of RSIS include a durable cell cycle arrest, initiated and maintained by the p16/p107/p130 axis and characterised by a panel of senescence-associated markers including activation of SASP factor. Whilst RSIS in the p16 positive cancer cell context appears to be independent of p53 or a DDR, it results in a nucleolar signature consistent with premature senescence. By contrast, RP knockdown in the p16-null BLBC cell line, MDA-MB-231, resulted in activation of the p53-caspase 9/3 cascade, suggesting a potentially divergent response dependent on p16-status or other factors.

Regulated and controlled ribosome biogenesis is critical for homeostasis, is fine-tuned to meet changing translational demands, and is intimately linked to the cell cycle. Cancer cells have an elevated translational requirement which may be required to support their dysregulated proliferative phenotype. Indeed, there is an existing link between RP silencing and cell cycle modulation, predominately in the context of p53-mediated mechanisms. For example, impairing ribosomal biogenesis in the cancer cell setting has been previously shown to lead to leakage of RPs, such as RPL11, followed by inhibition of MDM2 and a p53-mediated cell cycle arrest ^11, 13^. Depletion of specific RPs has also been shown in p16 negative cancer cell contexts to lead to supra-induction of p53 and cell cycle arrest ^11^. However, in mouse ESC/iPSCs, ribosomal stress leads to p53-dependent apoptosis ^14^, whilst depletion of RPL5/RPL11 in normal human lung fibroblasts led to a reduced ribosomal content and restrained cell cycle progression rather than a cell cycle arrest ^15^. The latter is in contrast to the RP hits identified here, whose knockdown failed to perturb cell cycle progression in a panel of normal fibroblasts and epithelial cells. Moreover, the clinical utility of compounds which induce nucleolar disruption is limited, as these agents can possess genotoxic activity leading to longer-term mutagenic effects. Conscious of this, we demonstrate that knockdown of our RPs hits did not alter nucleolar phenotype, integrity or abundance in normal HMECs, and generated an altered yet intact nucleolar morphology in the cancer cell setting in the absence of a DDR. More broadly, our observations suggest that nucleolar morphology represents a novel marker capable of distinguishing normal HMECs cellular senescence from premature senescence in HMECs or following cancer cell-specific RSIS. In support of this, Lessard et al. recently demonstrated that senescence-associated ribosome biogenesis defects contribute to cell cycle arrest through the Rb pathway which is associated with altered nucleolar morphology ^16^. In summary, we show that RP knockdown initiates a p16-mediated senescence programme in a p53-independent and cancer-specific manner, and paves the way for pro-senescence therapies aimed at re-engaging this potent tumour suppressor mechanism in highly aggressive breast cancer subtypes, such as BLBC.

In line with the findings that our RPs hits are dysregulated in both BLBC and p16^HIGH^ tumours, multiple RPs have been found to be overexpressed in a wide variety of human malignancies ^17, 18^. Further, ribosomeopathies, a collection of rare diseases characterised by genetic RP mutations, are often associated with an increased cancer risk, including breast cancer, indicating that disrupted ribosomal biosynthesis may play a role in cellular transformation ^19,20^. Of note, RPS7 missense mutations have been identified in 50-60% of patients with the cancer susceptibility syndrome, Diamond Blackfan anaemia. It has also been suggested that dysregulation of RPs supports the hyperproliferative cancer cell phenotype ^21^. This may result in an altered RP composition which favours the cancer translatome ^22^. Consequently, efforts are already underway to identify therapeutic agents which target the cancer ribosome. For example, compounds which target RNA Pol I and inhibit rRNA transcription, such as CX-3543 and CX-5461, are emerging as effective pro-apoptotic cancer agents ^23^. CX-3543 induces apoptosis in a panel of cancer cell lines, regardless of p53 status, and slows tumour growth within breast (MDA-MB-231) and pancreatic cancer xenograft models ^24^. The success of these compounds suggests that tumour cell addiction to enhanced ribosomal biosynthesis could be exploited for therapeutic gain, and our findings indicate that this could be achieved through a RSIS pro-senescence strategy. Both our work and the findings of Lessard et al. ^16^ indicate that a small molecule which perturbs ribosomal protein homeostasis could be effective at selectively engaging a p16-dependent senescence programme in cancer cells. In support of this, we present evidence of co-ordinated dysregulation of our two most potent RP hits, RPS3A and RPS7. The combination of these two RPs synergises with p16 leading to an earlier disease onset, and a poor patient outcome. As such, future endeavours might explore the utility of RPS3A and RPS7 as biomarkers. When combined with p16 (as presented here for breast tissue microarrays) and/or nucleolar morphological analysis this may enable better patient risk stratification and alternative personalised approaches for breast cancers. We propose cancer cell specific senescence activation via RSIS as a novel pro-senescence strategy which raises many future avenues for research. There is now compelling evidence that TIS cells are retained for many years following cytotoxic regimes, potentially pre-disposing cancer survivors to a multitude of age-related pathologies ^25-27^. Therefore, any future pro-senescence strategies would need form part of a combinatorial strategy. This may take the form of engaging immunosurveillance mechanisms following RSIS initiation, and provides a rationale for seeking novel senescent cancer cell ‘senolytics’ which could form part of a combinatorial therapeutic regime.

## ACKNOWLEDGEMENTS

MM was funded by the MRC (MR/K501372/1) and Barts Charity. MRS and JCG were supported by the Office of Health and Biological Research, US. Department of Energy under Contract No. DE-AC02-05CH11231.

## AUTHOR CONTRIBUTIONS

MM developed the hypothesis, designed the experimental approach, performed experiments, analysed the data, and wrote the manuscript. LG, SD and LJ contributed to the breast cancer tissue microarray generation and image analysis. JCG, MRS and JK contributed to writing the manuscript. MP contributed to experimental design and data interpretation. CLB conceived the hypothesis, led the project, designed the experimental approach assisted with experiments, interpreted data and wrote the manuscript.

